# Genetic colocalization atlas points to common regulatory sites and genes for hematopoietic traits and hematopoietic contributions to disease phenotypes

**DOI:** 10.1101/787333

**Authors:** CS Thom, BF Voight

## Abstract

**Background:** Genetic associations link hematopoietic traits and disease end-points, but most causal variants and genes underlying these relationships are unknown. Here, we used genetic colocalization to nominate loci and genes related to shared genetic signal for hematopoietic, cardiovascular, autoimmune, neuropsychiatric and cancer phenotypes.

**Results:** Our findings recapitulate developmental hematopoietic lineage relationships, identify loci associating traits with causal genetic relationships, and reveal novel associations. Out of 2706 loci with genome-wide significant signal for at least 1 blood trait, we identified 1779 unique sites (66%) with shared genetic signal for 2+ hematologic traits at a false discovery rate <5%. We could assign some sites to specific developmental cell types during hematopoiesis based on affected traits, including those likely to impact hematopoietic progenitor cells and/or megakaryocyte-erythroid progenitor cells. Through an expanded analysis of 70 human traits, we define 2+ colocalizing traits at 2007 loci from an analysis of 9852 sites (20%) containing genome-wide significant signal for at least 1 GWAS trait. In addition to variants and genes underlying shared genetic signal between blood traits and disease phenotypes that had been previously related through mendelian randomization studies, we define loci and related genes underlying shared signal between eosinophil count and eczema. We also identified colocalizing signals in a number of clinically relevant coding mutations, including in sites linking *PTPN22* with Crohns disease, *NIPA* with coronary artery disease and platelet trait variation, and the hemochromatosis gene *HFE* with altered lipid levels. Finally, we anticipate potential off-target effects on blood traits related novel therapeutic targets, including *TRAIL*.

**Conclusions:** Our findings provide a road map for gene validation experiments and novel therapeutics related to hematopoietic development, and offer a rationale for pleiotropic interactions between hematopoietic loci and disease end-points.

## Background

Identifying causal loci and genes from human genetic data is integral to elucidating novel disease insights and therapies approaches. Quantitative hematopoietic traits are relatively well studied, although relatively few causal variants and genes have been elucidated [1, 2]. Mendelian randomization studies have established causal relationships between hematopoietic traits and cardiovascular, autoimmune and neuropsychiatric disease [2]. Still, causal genes and loci remain elusive.

Genetic colocalization analysis permits identification of shared regulatory loci, with advances extending the scope of potential studies from two to >10 traits undergoing simultaneous analysis [3–5]. Recently, a colocalization algorithm was used to identify known and novel loci related to cardiovascular traits [5]. We reasoned that a similar analytical pipeline could help explain variants and genes underlying hematopoietic and other disease phenotypes. In this way, aggregated summary statistics might be used to specifically target loci with pleiotropic effects on multiple traits, enacted through one or a handful of genes.

Developmental cell types during hematopoiesis, the process that gives rise to all blood lineages, are relatively well mapped. We hypothesized that shared genetic signal impacting traits from multiple blood lineages might nominate genomic loci related to the stem and progenitor cells that spawned those types of blood cells. This approach is orthogonal to prior data that analyzed patterns in accessible chromatin to define genomic locations affecting multiple blood lineages [1]. For example, a shared single nucleotide polymorphism (SNP) related to quantitative variation in platelet, red blood cell (RBC), and white blood cell (WBC) counts might indicate a site or mechanism that is active in hematopoietic stem and progenitor cells (HSCs). SNPs related to platelet and RBC counts, but not WBC count, might reveal loci and related genes for megakaryocyte-erythroid progenitor (MEP) cells. We hypothesized that the directionality of such relationships might help elucidate lineage decisions during hematopoiesis, and help target loci and genes related to developmental hematopoiesis.

Blood traits are related to a number of human disease phenotypes [2]. Blood cells can cause disease (e.g. autoimmune traits) or be affected by therapies (e.g. anemia secondary to chemotherapy). For this reason, understanding pleiotropic associations between blood and other traits could reveal translationally relevant trait relationships or help predict off-target effects of gene-modifying therapies.

Here, we used genetic colocalization to define sites wherein 2 or more human traits shared genetic signal at genome-wide significant loci. We initially examined blood traits, and later expanded our analysis to include a total of 70 blood, autoimmune, cardiovascular, cancer, and neuropsychiatric traits. We then looked for quantitative trait loci impacting gene expression (eQTL) or exon splicing variation (sQTL) at or near sites of genetic colocalization. Our results identify sites that affect specific cell types during hematopoietic development, and reveal genetic variants underlying trait relationships between blood parameters and disease end-points.

## Results

### Genetic colocalization recapitulates hematopoietic lineage relationships

Our first aim was to validate whether colocalization could effectively capture known trait relationships and genetic correlations between hematopoietic lineages [1]. We performed colocalization analysis [5] using summary statistics related to 34 quantitative hematopoietic traits for 2706 genome-wide significant loci [2], revealing a total of 1779 sites wherein 2 or more traits colocalized with a posterior probability > 0.7 (**Additional file 1: Table S1**). In simulations, these criteria identified the causal variant, or a variant in high LD with the causal variant, with a false discovery rate < 5% [5]. Colocalization sites specified 3.6±2.3 traits (mean±SD), with 22% of the loci (259 in total) representing highly pleiotropic sites where 6 or more traits colocalized (**Fig. 1a**). Hence, a substantial proportion of interrogated loci (66%) impacted multiple hematopoietic traits.

**Figure 1.**
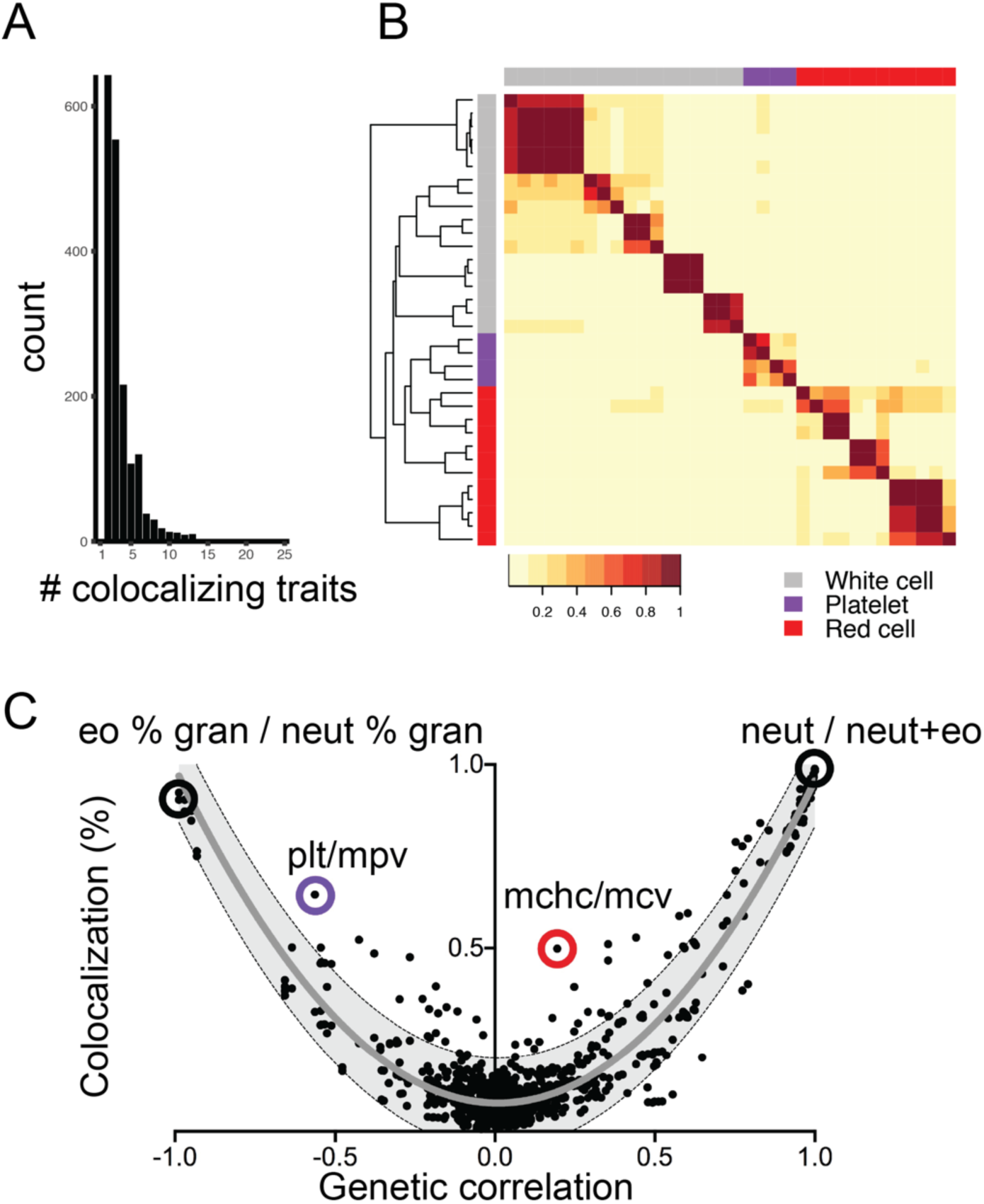
Genetic colocalization between blood traits reflects hematopoietic lineage relationships. **a** Number of traits identified at each colocalization site (max = 25). **b** Heat map depicting percent overlap at colocalization sites between each hematopoietic trait pair. In each box, the number of sites where the row-specified trait and column-specified trait colocalized was normalized to the total number of colocalization sites for the ‘row trait’. For this reason, the heat map is asymmetric. Color scale represents the proportion of loci where each pair of traits colocalized. To the left of the heat map, hierarchical clustering accurately segregated red cell, platelet, and white cell traits in general agreement with blood lineage relationships. **c** Degree of colocalization (% overlap) generally reflects genetic correlation between trait pairs. Shaded area depicts 95% prediction, with gray line at mean. Exemplary trait pairs are highlighted. eo % gran, percentage of granulocytes that are eosinophils. neut % gran, percentage of granulocytes that are neutrophils. plt, platelet count. mpv, mean platelet volume. mchc, mean corpuscular hemoglobin content. mcv, mean red cell corpuscular volume. neut, neutrophil count. neut+eo, total neutrophil plus eosinophil count.

To investigate trait relationships, we constructed a heat map to depict the percentage colocalization between trait pairs (**Fig. 1b**). Hierarchical clustering of colocalization results reflected blood lineage relationships, with platelet, erythroid, and white blood cell traits generally clustering as expected.

We then asked whether our colocalization findings mirrored genetic correlation between hematopoietic traits [6]. Indeed, more closely related traits colocalized more often (**Fig. 1c**, r^2^=0.91 by quadratic regression with least squares fit). Directly correlated (e.g., ‘neutrophil count’ and ‘neutrophil + eosinophil count’; ‘granulocyte count’ and ‘myeloid white blood cell count’), and inversely correlated trait pairs (e.g., ‘eosinophil percent of granulocytes’ and ‘neutrophil percent of granulocytes’; ‘lymphocyte percent’ and ‘neutrophil percent’), essentially always colocalized. Several trait pairs fell outside the 95% prediction interval (e.g. ‘mean platelet volume’ and ‘platelet count’; ‘mean corpuscular hemoglobin concentration’ and ‘mean red cell volume’) (**Fig. 1c**). Lineage-critical loci or genes might be expected to have more significant influence on these lineage-restricted trait pairs than would be captured by genetic correlation measure.

In sum, these results validated the notion that colocalization analysis results would mirror genetic correlation, and reflect known relationships among hematopoietic lineages and traits. Interestingly, trait pairs without genetic correlation frequently had some degree of colocalization (**Fig. 1c**, y-intercept = 0.077±0.123). This likely reflects horizontal pleiotropy, in which a given locus and related gene(s) impact traits that are not biologically related. In the context of hematopoietic development, our derived estimate of chance colocalization between unrelated traits is therefore ∼8%.

### A genetic colocalization strategy to identify loci related to hematopoietic development

We then leveraged our colocalization results to identify quantitative trait loci (QTLs) related to specific hematopoietic lineages and cell types. For example, loci where white blood cell (WBC), red blood cell (RBC), and platelet counts colocalize might indicate developmental perturbation in hematopoietic stem and progenitor cells (HSCs). We therefore looked for sites of colocalization between these quantitative blood traits, and identified overlapping genome-wide significant QTLs. Indeed, QTLs related to these loci pointed to known HSC regulatory genes *SH2B3* [7, 8], *ATM* [9], and *HBS1L-MYB* [10] (**Additional file 1: Table S2**).

We also parsed loci identified by colocalization to specifically affect platelet or red cell traits, with the hypothesis that these loci would relate to terminally differentiated blood cell biology. There were 439 sites nominated by colocalization analysis specifically for red cell traits (RBC, HCT, MCV, MCH, MCHC, RDW) but *not* platelet traits or WBC count. These sites, or highly linked loci, influenced expression of 614 genes (123 genes in whole blood, **Additional file 1: Table S3**). Gene ontology (GO) analysis [11] of these gene set revealed significant enrichment of genes related to cellular metabolic processes (**Additional file 1: Table S4**). A similar analysis of platelet trait-restricted sites (PLT, PCT, MPV, PDW), including highly linked loci, identified 270 sites impacting expression of 399 genes (77 genes in whole blood, **Additional file 1: Table S5**). Pathway analysis of these genes revealed enrichment of apoptotic cell clearance and metabolic processes (**Additional file 1: Table S6**). Complement-mediated apoptotic cell clearance mechanisms are indeed important for regulating platelet count [12].

To our surprise, pathways analyses of red cell and platelet lineage-restricted colocalization QTLs were not enriched processes ascribed to hematopoiesis, erythropoiesis, or megakaryopoiesis. This suggests that genes and processes linked to terminal red cell and platelet traits are largely impacted by cellular function and reactivity, rather than developmental perturbations. With notable exceptions whereby causal loci do impact hematopoietic development (e.g., [7, 13–15]), our findings suggest that this may not be true for the majority of identified genes and factors. In fact, our results indicate that cell-extrinsic properties (e.g. apoptotic cell clearance mechanisms) frequently impact quantitative hematopoietic traits. In sum, our findings reveal a multitude of known variants and genes, as well as novel QTL and related genes that warrant further study.

### Illuminating hematopoietic contributions and associations with disease phenotypes

We then applied an extended colocalization analysis to summary statistics for 70 total hematopoietic, cardiovascular, autoimmune, cancer, and neuropsychiatric traits (**Additional file 1: Table S7**). Following allele harmonization, colocalization analysis using 9852 genome-wide significant loci from the NHGRI database [16] and blood traits [2] revealed a total of 2007 sites (20%) wherein 2 or more traits colocalized with a posterior probability > 0.7 (**Additional file 1: Table S8**). The average number of traits that colocalized at a given site was 3.3±2.4 (mean±SD), with 84 loci identified as a ‘very pleiotropic’ colocalization site for ≥9 traits (**Fig. 2a**). Known trait relationships were recapitulated among these colocalization sites (e.g. bipolar disorder and schizophrenia; **Fig. 2b-d**). These results again reflected genetic correlation between traits, estimating a small degree of pleiotropy (∼4%) absent genetic correlation (**Fig. 2d**, r^2^=0.86, y-intercept= 0.041±0.120).

**Figure 2.**
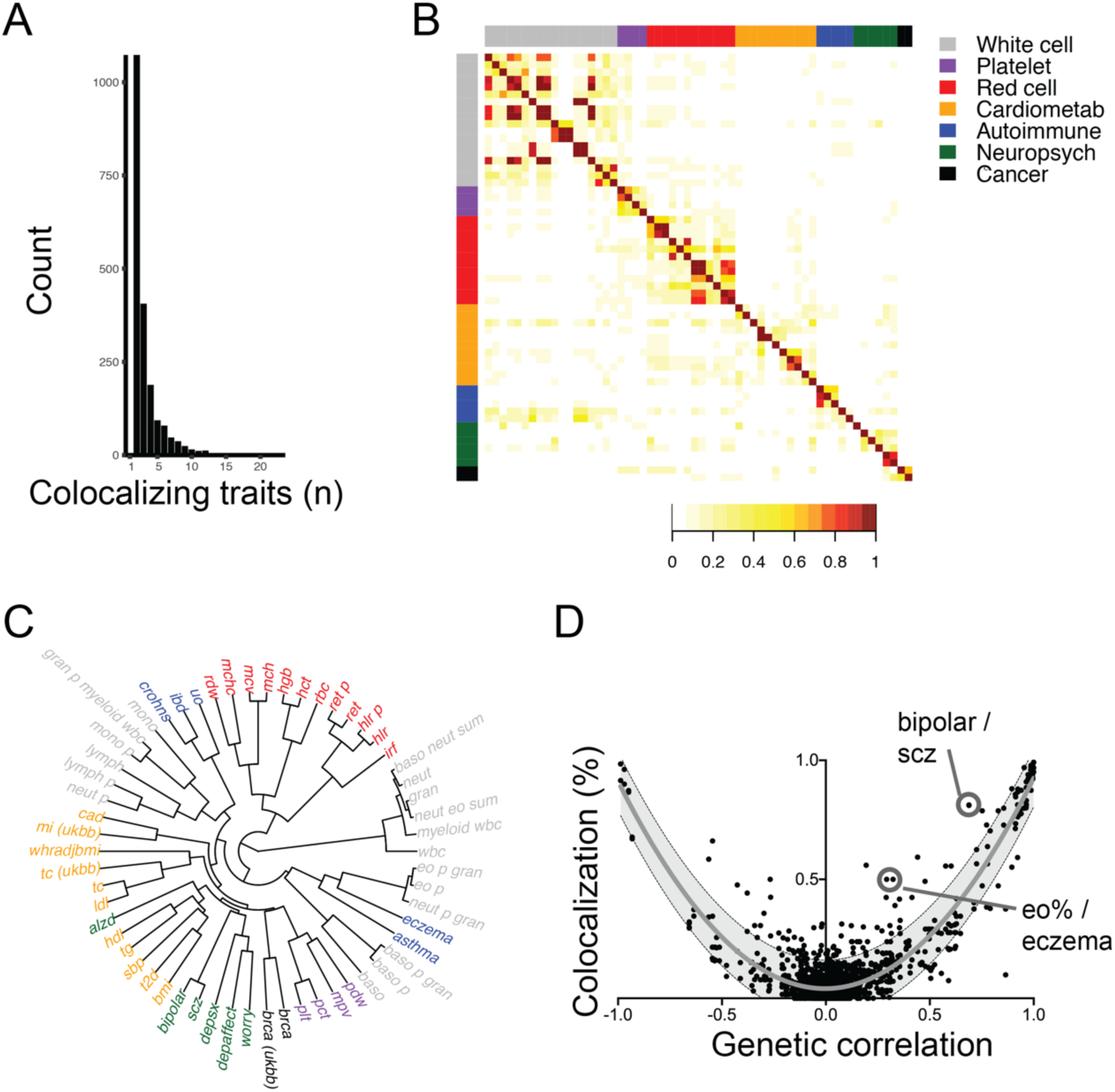
Genetic colocalization reveals shared regulatory loci and implicates causal genes underlying genetic associations between hematopoietic traits and disease end-points. **a** Number of traits identified at each colocalization site (max = 23). **b** Heat map depicting percent overlap at colocalization sites between each trait pair. In each box, the number of sites where the row-specified trait and column-specified trait colocalized was normalized to the total number of colocalization sites for the ‘row trait’. For this reason, the heat map is asymmetric. **c** Hierarchical clustering based on colocalization results accurately associates related traits, which are color coded according to the key in part **b. d** Degree of colocalization (% overlap) reflects genetic correlation between trait pairs. Shaded area depicts 95% prediction, with gray line at mean. Exemplary trait pairs are circled. scz, schizophrenia. eo%, percentage of white blood cells that are eosinophils.

Mendelian randomization analyses have established causal relationships between blood traits and some disease phenotypes [2]. Despite holding significant therapeutic potential, most causal loci and genes underlying these associations are unknown. Our results reveal putatively causal loci, related genes and molecular pathways related to these trait pairs (**Additional file 1: Table S9-S16**). For example, whole blood QTLs related to genes known to affect asthma pathogenesis or severity (e.g. *IL18R* [17–19], *ZFP57* [20], *BTN3A2* [21], *NDFIP1* [22], *SMAD3* [23], *CLEC16A* [24], and *TSLP* [25]) were associated with colocalization sites for asthma and neutrophil, eosinophil, monocyte and/or lymphocyte traits (**Additional file 1: Table S9-S12**). Similarly, QTLs for genes linked to coronary artery disease risk (e.g. *XRCC3* [26], *SREBF1* [27], *GIT1* [28, 29], *SKIV2L* [29], *MAP3K11/MLK3* [30]) were associated with colocalization sites linking coronary artery disease with lymphocyte and/or reticulocyte counts (**Additional file 1: Table S13-S15**). Other identified genes associated with colocalization loci represent novel findings that could enhance understanding of the pathophysiology and/or treatment of these diseases, although functional validation remains necessary.

Our findings also revealed novel trait associations. Although eosinophil percentage and eczema did not reach statistical significance by Mendelian Randomization, these traits are clinically related [31] and colocalized at 10 sites (**Additional file 1: Table S16**). These results were localized near genes that regulate eosinophil biology (*ETS1* [32, 33] and *ID2* [33]) and autoimmune disease (*KIAA1109* [34–36] and *TAGAP* [37]), and indicate potential regulation of unexpected genes that warrant validation (*SNX32, ZNF652, KLC2*).

### Colocalization at coding variation sites identifies clinically relevant trait associations

Next, we reasoned that colocalizing sites could help explain unexpected or pleiotropic effects of gene perturbations. Here, we focused on missense variation in coding regions to establish direct locus-gene relationships. This approach identified clinically relevant cross-trait associations.

Variation in rs2476601 causes a missense mutation in *PTPN22* (Cys1858Thr). This site has been linked to autoimmunity and Crohns disease phenotypes, but not ulcerative colitis [38]. Immune response dysregulation, including WBC biology, contributes to the Crohns phenotype [38]. We identified shared genetic signal for increased Crohns disease risk and decreased WBC count, but not ulcerative colitis, at this location (**Additional file 1: Table S17**). This finding supports a specific clinical association with Crohns for the *PTPN22* Cys1858Thr mutation.

Mean platelet volume (MPV) variation has previously been linked to altered risk of coronary artery disease, but understanding of genes underlying this association is lacking [2]. We identified colocalizing signals for increased coronary artery disease risk and increased MPV in a missense coding mutation for *ZC3HC1/NIPA* (**Additional file 1: Table S13**). This variant causes an Arg>His missense change in several *NIPA* isoforms. *NIPA* impacts heart disease risk and cell cycle regulation [39]. Further studies are needed to understand how this gene might coordinately impact platelet biology and coronary artery disease risk, as well as other traits linked to this locus.

Altered lipid and cholesterol levels have been clinically observed in patients with hereditary hemochromatosis due to mutations in *Human Factors Engineering* (*HFE*) [40]. Patients with hemochromatosis have lower cholesterol levels than normal, although an open question is whether this observation is due to manifestations of disease or *HFE* deficiency itself. Our data show that individuals heterozygous for the Cys282Tyr allele have lower reticulocyte count and higher total cholesterol and low density lipoprotein levels (**Additional file 1: Table S18**). This suggests that *HFE* haploinsufficiency increases cholesterol and lipid levels, and that decreased cholesterol in patients with hemochromatosis occurs secondary to myriad tissue manifestations of clinically significant hemochromatosis or iron overload [41].

Finally, we hypothesized that our analysis might also help predict off-target effects of novel therapeutic agents. For example, tumor necrosis factor (TNF)-related apoptosis inducing ligand (*TRAIL*) is a promising novel chemotherapeutic target [42]. A mutation in the *TRAIL* 3’ UTR was associated with decreased total cholesterol and triglyceride levels, as well as decreased platelet counts (**Additional file 1: Table S19**). It will be interesting to see whether these traits are affected in upcoming clinical trials targeting *TRAIL*.

## Discussion

Genetic colocalization approaches have proven a powerful tool in revealing pleiotropic effects of certain loci on multiple traits [3, 4]. Here, we have adapted the colocalization methodology to reveal sites and genes related to specific cell stages in hematopoietic development, and identify relevant trait relationships between blood traits and human disease end-points. We present what we believe to be a minimal estimate of these associations, given our conservative threshold for colocalization (PP>0.7). This threshold revealed high-confidence targets, although future gene discovery studies might instead use a more relaxed threshold (e.g., PP>0.5) to enable a more encompassing set of loci.

GWAS have linked thousands of genomic sites with blood trait variation [2]. The biology related to each site could relate to developmental hematopoiesis, as has been shown for *CCND3* [13], *CCNA2* [14], *SH2B3* [7], and *RBM38* [15]. Alternatively, biology related to GWAS sites might impact terminally differentiated cell reactivity or turnover. For example, altered platelet reactivity can affect quantitative platelet traits [43, 44]. Cellular validation experiments might be streamlined if one could better parse relevant sites, genes and developmental stages based on GWAS information. Gene targets presented herein represent one approach to such a computational pipeline, and are orthogonal to previously published findings based on accessible chromatin patterns during hematopoietic development [1]. Future studies combining these computational modalities might be useful for those interested in evaluating specific genes or loci in blood progenitor biology.

Our expanded analysis of 70 human traits recapitulated known trait relationships between blood traits and human disease phenotypes, and identified sites with potential translational relevance. Understanding how missense coding mutations impact phenotypes offers the most direct relationship between genes and traits. An adaptation of our colocalization strategy might be employed to predict off-target effects of gene modulation, help understand cellular basis of disease, or investigate unexpected cellular developmental relationships (e.g., sites related to multiple mesoderm-derived tissues might triangulate to early mesodermal biology). We anticipate an expanded array of such targets could be revealed with larger, trans-ethnic GWAS.

## Conclusion

In an extensive genetic colocalization analysis, we have identified loci, genes and related pathways related to hematopoietic development. Further, our colocalization results identified loci relating 70 hematopoietic, cardiovascular, autoimmune, neuropsychiatric and cancer phenotypes. This repository of associations will be useful for mechanistic studies aimed at understanding biological links between phenotypes, for developing novel therapeutic strategies, and for predicting off-target effects of small molecule and gene therapies.

## Methods

### SNP and study selection

Human genome version hg19 was used for all analyses. GWAS summary statistics were obtained from publicly available repositories (**Additional file 1: Table S7**). We narrowed analysis to just those GWAS summary statistics for European populations with >1 x10^6^ sites (i.e., those that were genome-wide). Analyzed SNPs were identified as genome-wide significant in the largest hematopoietic trait GWAS to date [2] or from a repository of genome-wide significant SNPs from a compilation of GWAS from the NHGRI [16] (downloaded January 2019). In addition, we analyzed quantitative trait locus data from GTEx V7 [45].

### Colocalization analyses

We used HyPrColoc software to conduct colocalization experiments [5]. This software requires effect (e.g., beta or odds ratio) and standard error values for each analyzed SNP. We chose to analyze based on chromosome and position, given that multiple rsIDs might overlap at a given locus and be inconsistent between different GWAS. Although this removed duplicate rsIDs and may have caused some bias, we reasoned that this would be a minority of sites. This strategy optimized the number of individual positions that we were able to incorporate into our input dataset for colocalization analysis. We specifically looked at 500 kb regions (250 kb on either side of each site), in line with prior colocalization literature [5].

As the SNPs considered as input data varied between analyses, we presented separate results from analysis of 34 hematologic traits, and a composite 70 traits. GWAS summary statistics were harmonized prior to analyses (https://github.com/hakyimlab/summary-gwas-imputation/wiki). There were 29,239,447 genomic sites analyzed for colocalization among the hematologic traits. A total of 1,544,623 harmonized sites were analyzed from GWAS summary statistics for the 70 traits. The decreased number of sites included in this latter analysis resulted in decreased power to detect associations. This was reflected in the maximum number of traits colocalized in **Fig. 1a** was 25 (out of 34 traits) versus 23 traits in **Fig. 2a** (out of 70 traits).

After colocalization analysis, we narrowed our focus on only those loci with posterior probability for colocalization (PPFC) >0.7 based on empiric simulations results from the creators of this algorithm showing that this conservatively gave a false discovery rate <5% [5]. We note that a more relaxed PPFC (e.g. >0.5) yields substantially more loci. A less conservative threshold could in this way be used as a hypothesis-generating experiment for cellular follow up studies.

### Coding variant identification

We used the Ensembl Variant Effect Predictor (http://grch37.ensembl.org/Homo_sapiens/Tools/VEP) to identify coding variants and related gene consequences.

### Linkage disequilibrium and quantitative trait locus (QTL) analyses

We wanted to assess comprehensively the potential gene expression or splicing changes related to colocalization sites. Thus, we analyzed each colocalization site together with all sites in high linkage disequilibrium (EUR r^2^>0.90, PLINK version 1.9).

We used closestBed (https://bedtools.readthedocs.io) to identify the nearest gene to each SNP. Genes and positions were defined by BioMart (http://www.biomart.org/).

For each group of linked SNPs around a colocalization locus, we identified all eQTLs (GTEx V7 [45]), as well as all sQTLs as defined by two different algorithms (sQTLseekeR [46], Altrans [47]). In the manuscript and Additional file 1, the quantity of QTL SNPs and pathway analyses reflect a composite of all genes impacted by a given locus, or by highly linked SNPs. Note that a given colocalization site might be linked with several SNPs, and that these SNPs might be in proximity and/or impact different genes. Affected genes shown are those with a unique Ensembl gene identifier (ENSG). In some cases, gene names may differ between Nearest Gene, eQTL and sQTL columns given that the underlying analyses were derived from different catalogues.

### Gene ontology analysis

We submitted QTLs associated with specific traits for biological pathway assessment using the Gene Ontology (GO) resource (http://geneontology.org/). Statistical significance of GO Biological Process enrichments were assessed using binomial tests and Bonferroni correction for multiple testing. Presented data were those pathways with p<0.05.

### Empirical distribution for expected colocalization counts

We used LDSC to estimate genetic correlation between traits (v1.0.1) [6]. Presented genetic correlation data reflect rg values obtained from LDSC analysis.

## Data presentation

Data were created and presented using R, LocusZoom (http://locuszoom.org), Adobe Illustrator CS6 and GraphPad Prism 8.

## Statistics

Statistical analyses were conducted using R and GraphPad Prism 8.

## Declarations

### Ethics approval and consent to participate

Consent was obtained as part of original genetic studies. Hence, no further consent was necessary to use summary statistics as part of our study.

### Consent for publication

Not applicable

### Availability of data and materials

Code and scripts used to generate and analyze data are available on GitHub (https://github.com/thomchr/2019.Coloc).

### Authors’ contributions

CST and BFV conceived of the project, generated, analyzed and interpreted data, and wrote the paper.

## Acknowledgements

Not applicable

## Additional file 1 Table Legends

**Table S1.** Traits and SNPs identified by colocalization analysis [5] of hematopoietic traits. All identified sites are shown in this table. Candidate SNPs are indicated as chr:pos. The posterior probability of colocalization, regional (genomic) probability of colocalization, and posterior probability explained at each locus are indicated.

**Table S2.** ‘Hematopoietic stem cell’ sites at which white blood cell (wbc), red blood cell (rbc), and platelet (plt) counts colocalize. Sites were specified by chromosome and position. The rsID(s) associated with each site are shown. Gene symbols for the nearest gene, all eQTLs, all eQTLs in whole blood, and all sQTLs (sQTLseekeR and Altrans methods) are shown in the indicated columns related to the indicated SNP (rsID) and any SNPs in high linkage disequilibrium (r^2^>0.9).

**Table S3.** ‘RBC trait only’ sites at which only the indicated red blood cell traits colocalized, excluding platelet or white blood cell traits. Gene symbols for the nearest gene, all eQTLs, all eQTLs in whole blood, and all sQTLs (sQTLseekeR and Altrans methods) are shown in the indicated columns related to the indicated SNP (rsID) and any SNPs in high linkage disequilibrium (r^2^>0.9). rbc, red blood cell count. hct, hematocrit. mcv, mean red cell corpuscular volume. rdw, red cell distribution width.

**Table S4.** Gene ontology pathway analysis of genes regulated by eQTLs linked to ‘RBC trait only’ sites. Shown are pathways with p<0.05 by Binomial test using Bonferroni correction for multiple testing.

**Table S5.** ‘Platelet trait only’ sites at which only the indicated platelet traits colocalized, excluding red blood cell or white blood cell traits. Gene symbols for the nearest gene, all eQTLs, all eQTLs in whole blood, and all sQTLs (sQTLseekeR and Altrans methods) are shown in the indicated columns related to the indicated SNP (rsID) and any SNPs in high linkage disequilibrium (r^2^>0.9). plt, platelet count. pct, platelet-crit. mpv, mean platelet volume. pdw, platelet distribution width.

**Table S6.** Gene ontology pathway analysis of genes regulated by eQTLs linked to ‘platelet trait only’ sites. Shown are pathways with p<0.05 by Binomial test using Bonferroni correction for multiple testing.

**Table S7.** Genome wide association study summary statistics used in our analysis. The trait(s) queried are shown, along with study Pubmed identification number (PMID). UK BioBank studies can be found using the link provided.

**Table S8.** Traits and SNPs identified by colocalization analysis [5] of 70 human traits. All identified sites are shown in this table. Candidate SNPs are indicated as chr:pos. The posterior probability of colocalization, regional (genomic) probability of colocalization, and posterior probability explained at each locus are indicated.

**Table S9.** Colocalization sites for Lymphocyte count (lymph) and Asthma. Gene symbols for the nearest gene, all eQTLs, all eQTLs in whole blood, and all sQTLs (sQTLseekeR and Altrans methods) are shown in the indicated columns related to the indicated SNP (rsID) and any SNPs in high linkage disequilibrium (r^2^>0.9).

**Table S10.** Colocalization sites for neutrophil count (neut) and Asthma. Gene symbols for the nearest gene, all eQTLs, all eQTLs in whole blood, and all sQTLs (sQTLseekeR and Altrans methods) are shown in the indicated columns related to the indicated SNP (rsID) and any SNPs in high linkage disequilibrium (r^2^>0.9).

**Table S11.** Colocalization sites for eosinophil percentage (eo%) and Asthma. Gene symbols for the nearest gene, all eQTLs, all eQTLs in whole blood, and all sQTLs (sQTLseekeR and Altrans methods) are shown in the indicated columns related to the indicated SNP (rsID) and any SNPs in high linkage disequilibrium (r^2^>0.9).

**Table S12.** Colocalization sites for monocyte count (mono) and Asthma. Gene symbols for the nearest gene, all eQTLs, all eQTLs in whole blood, and all sQTLs (sQTLseekeR and Altrans methods) are shown in the indicated columns related to the indicated SNP (rsID) and any SNPs in high linkage disequilibrium (r^2^>0.9).

**Table S13.** Colocalization sites for mean platelet volume (mpv) and coronary artery disease (cad). Gene symbols for the nearest gene, all eQTLs, all eQTLs in whole blood, and all sQTLs (sQTLseekeR and Altrans methods) are shown in the indicated columns related to the indicated SNP (rsID) and any SNPs in high linkage disequilibrium (r^2^>0.9).

**Table S14.** Colocalization sites for reticulocyte count (ret) and coronary artery disease (cad). Gene symbols for the nearest gene, all eQTLs, all eQTLs in whole blood, and all sQTLs (sQTLseekeR and Altrans methods) are shown in the indicated columns related to the indicated SNP (rsID) and any SNPs in high linkage disequilibrium (r^2^>0.9).

**Table S15.** Colocalization sites for lymphocyte count (lymph) and coronary artery disease (cad). Gene symbols for the nearest gene, all eQTLs, all eQTLs in whole blood, and all sQTLs (sQTLseekeR and Altrans methods) are shown in the indicated columns related to the indicated SNP (rsID) and any SNPs in high linkage disequilibrium (r^2^>0.9).

**Table S16.** Colocalization sites for eosinophil percentage (eo%) and Eczema. Gene symbols for the nearest gene, all eQTLs, all eQTLs in whole blood, and all sQTLs (sQTLseekeR and Altrans methods) are shown in the indicated columns related to the indicated SNP (rsID) and any SNPs in high linkage disequilibrium (r^2^>0.9).

**Table S17.** Analysis of a coding variant (rs2476601) that causes a missense mutation in *PTPN22* shows significant colocalization white white blood cell (wbc) count and Crohns disease. The effect sizes and direction (+/-) are shown.

**Table S18.** Colocalization analysis for a coding variant (rs1800562) in Human Factors Engineering (*HFE*), mutations in which cause hereditary hemochromatosis. Effects on total cholesterol (TC), low density lipoprotein (LDL), and red blood cell traits (high light scatter reticulocyte count, hlr; high light scatter reticulocyte percentage, hlr_p; mean corpuscular hemoglobin concentration, mchc; red cell distribution width, rdw; reticulocyte count, ret; reticulocyte percentage, ret_p), with significant colocalization signal at this locus, are shown.

**Table S19.** Colocalization analysis for a coding variant (rs17600346) in Tumor necrosis factor (TNF)-related apoptosis inducing ligand (*TRAIL*, also known as *TNF10*). Effects on colocalized traits total cholesterol (TC), triglyceride (TG), white blood cell (granulocyte percentage of myeloid white blood cells, gran_p_myeloid_wbc; monocyte percentage, mono_p) and platelet traits (platelet-crit, pct; platelet count, plt) are shown.

